# Multi-trait meta-analyses reveal 25 quantitative trait loci for economically important traits in Brown Swiss cattle

**DOI:** 10.1101/517276

**Authors:** Zih-Hua Fang, Hubert Pausch

## Abstract

**Background:** The Brown Swiss dual-purpose cattle breed is renowned for high milk and protein yield and long productive lifetime under different production conditions. However, little is known about the genetic architecture of those traits because only few genome-wide association studies (GWAS) have been carried out in this breed. Moreover, most GWAS had been performed for single traits, thus preventing insights into potentially existing pleiotropic effects of trait-associated loci.

**Results:** To compile a comprehensive catalogue of large-effect QTL segregating in Brown Swiss cattle, we carried out association tests between partially imputed genotypes at 598,016 SNPs and daughter-derived phenotypes for more than 50 economically important traits, including milk production, growth and carcass quality, body conformation, reproduction and calving traits in 4,578 artificial insemination bulls from two cohorts of Brown Swiss cattle (Austrian-German and Swiss populations). Across-cohort multi-trait meta-analyses of the results from the single-trait GWAS revealed 25 quantitative trait loci (QTL; *P* < 8.36 x 10^−8^) for economically relevant traits on 17 *Bos taurus* autosomes (BTA). Evidence of pleiotropy was detected at five QTL located on BTA5, 6, 17, 21 and 25. Of these, two QTL at BTA6:90,486,780 and BTA25:1,455,150 affect a diverse range of economically important traits, including traits related to body conformation, calving, longevity and milking speed. Furthermore, the QTL at BTA6:90,486,780 seems to be a target of ongoing selection as evidenced by an integrated haplotype score of 2.49 and significant changes in allele frequency over the past 25 years, whereas either no or only weak evidence of selection was detected at all other QTL.

**Conclusions:** Our findings provide a comprehensive overview of QTL segregating in Brown Swiss cattle. Detected QTL explain between 2 and 10% of the variation in the daughter-derived phenotypes and thus may be considered as the most important QTL segregating in the Brown Swiss cattle breed. Multi-trait association testing boosts the power to detect pleiotropic QTL and assesses the full spectrum of phenotypes that are affected by trait-associated variants.

## Background

Genome-wide association studies (GWAS) between economically important traits and dense single nucleotide polymorphism (SNP) genotypes identified hundreds of quantitative trait loci (QTL) and thousands of trait-associated genetic variants in many cattle breeds [1]. Prioritizing trait-associated variants in genomic prediction models may accelerate genetic gain, particularly when large reference populations are not available, e.g., for traits that are either expensive or difficult to measure [2]. Furthermore, characterizing the spectrum of traits that are affected by pleiotropic QTL may contribute to a better understanding of the physiological underpinnings of economically important traits and help unravelling molecular-genetic mechanism through which QTL acts. This knowledge could further facilitate an improved prediction of correlated responses given selection for particular traits [3].

Natural or artificial selection changes the frequency of variants underpinning the traits under selection and their neighboring polymorphic sites in linkage disequilibrium. Strong selection may result in regions of extended homozygosity along the genome, i.e., selective sweeps (see review [4]). Detecting such patterns could provide insight into responses of the cattle genome to past and ongoing natural and artificial selection and reveal loci and variants that underpin adaptive and economically important traits [5], thus enhancing our understanding of genetic mechanisms controlling phenotypes under selection [6, 7].

The Brown Swiss cattle breed is popular in many European countries including Austria, Germany and Switzerland. Brown Swiss cows produce high milk and protein yields and have a long productive life under various production and environmental conditions. However, the genetic architecture of these traits is not well characterized in Brown Swiss cattle since only few GWAS were carried out so far in this breed [8–11]. Because most of the association studies performed so far considered either only one trait at a time or were restricted to clusters of related traits, detecting QTL that are associated with seemingly unrelated traits was not possible in those studies and the extent to which pleiotropic QTL contribute to trait variation and correlation in Brown Swiss cattle is currently unknown.

In this study, we performed GWAS for more than 50 economically important traits, including milk production, body conformation, carcass quality, and functional traits (reproduction, health and management), in Brown Swiss cattle. Our aim was to detect the most important QTL that underpin these economically important traits and investigate if the detected QTL showed pleiotropic effects or evidence of recent or ongoing selection.

## Methods

### Studied populations, phenotypes and genotypes

We considered genotypes and phenotypes of 4,578 Brown Swiss bulls that were used for artificial insemination from Austria, Germany and Switzerland. Estimated breeding values (EBV) and corresponding reliabilities for 56 economically important traits, including milk production, body conformation, carcass and meat quality, health and reproduction (Additional file 1: Table S1) were provided from the Swiss routine genetic evaluation for 1,875 bulls and the joint Austrian and German routine genetic evaluation [12] for 2,703 bulls. For our association analyses, we considered only bulls that had EBVs with reliabilities greater than 0.4 for all traits.

Of the 4,578 bulls, 870 bulls were genotyped with the Illumina HD Bovine SNP chip comprising 777,962 SNPs and 3,708 bulls were genotyped using the BovineSNP50 Bead chip (Illumina, San Diego, CA) comprising either 54,001 (version 1) or 54,609 (version 2) SNPs. The physical positions of the SNPs were determined based on the UMD3.1 assembly of the bovine genome [13]. Mitochondrial, Y-chromosomal SNPs and SNPs that had unknown chromosomal position were not considered for the subsequent analyses. Quality control including the exclusion of SNPs and animals with genotyping rates less than 80% and SNPs showing deviation from the Hardy-Weinberg equilibrium (*P* < 0.00001) was applied to the 50K and HD datasets separately using *plink* (version 1.9) software [14]. Following quality control, the 50K and HD datasets included 3,708 and 870 animals, respectively, that had genotypes at 42,649 and 647,091 SNPs. The 50K genotypes were imputed to higher density using a combination of *Eagle* (version 2.4) [15] and *Minimac3* (version 2.0.1) [16] as described in Pausch et al. [17]. Following the imputation, our final dataset consisted of 4,578 animals that had partially imputed genotypes at 642,196 SNPs. X-chromosomal (N=15,981) SNPs and 28,199 SNPs with minor allele frequency (MAF) less than 1% were not considered for further analyses, resulting in 598,016 SNPs that were considered for GWAS and selection signature analyses.

### Genome-wide association analyses

#### Single-trait association testing

For each trait considered, single-marker-based association studies were carried out using a two-step variance component-based approach that has been implemented using *emmax* software [18]: y = μ + u + e, where y is a vector of phenotypes, μ is the intercept, u is a vector of additive genetic effects 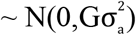, where 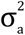 is the additive genetic variance and G is the genomic relationship matrix built using partly imputed HD genotypes at autosomal SNPs using *plink* (version 1.9) software [14]. e is a vector of residuals 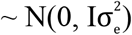, where I is the identity matrix and 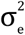 is the error variance. The allele substitution effect (b) was subsequently obtained from a generalized linear regression model: y = μ + Xb + η, where X is a vector of genotypes and η is a vector of random residual deviates with variance 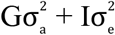.

#### Multi-trait meta-analysis

An approximate multi-trait test statistic was calculated for each SNP according to Bolormaa et al. [3] using 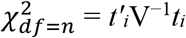, where n is the number of traits, *t*_*i*_ is a n x 1 vector of the signed t-values at the i^th^ SNP 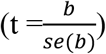 across traits and V^−1^ is the inverse of the n x n correlation matrix where the correlations were calculated over all signed t-values for each pair of traits.

SNPs with *P* value less than 8.36 x 10^−8^ (corresponding to a 5 % Bonferroni-corrected significance threshold) were considered as significantly associated for both single- and multi-trait analyses. QTL regions were defined as 5 million basepair (Mb) intervals centered on the most significantly associated (lead) SNP. The proportion of variance in daughter-derived phenotypes explained by a QTL was calculated according to Kemper et al. [19] using (*t*^2^−1)/N, where *t* is the t-value of the lead SNP at a given QTL for the trait under analysis, and N is the number of animals in the analysis.

### Detection of signatures of selection

Genomic regions showing evidence of selection were detected using either runs of homozygosity (ROH) or the integrated haplotype score (iHS) test. To investigate if the detected QTL were under selection, the extended haplotype homozygosity (EHH) was calculated surrounding the lead SNPs of these QTL. To assess changes in QTL frequency over time, allele frequency of the lead SNPs was calculated for the past 25 years based on the birth year of the studied Brown Swiss bulls.

We considered ROH with minimum length of 2 Mb, which are supposed to originate from common ancestors for up to 10-25 generations ago. This corresponds to approximately 50-150 years assuming an average generation interval of 5-6 years in cattle, to explore the impacts of recent artificial selection for economically important traits [20, 21]. ROHs that contained at least 50 SNPs and were longer than 2 Mb were identified using a sliding window-based approach implemented in *plink* (version 1.9) software [14]. Briefly, a sliding window consisting of 50 adjacent SNPs was shifted along all autosomes in steps of one SNP. Two missing genotypes and one heterozygous genotype were allowed within each sliding window. The minimum SNP density within a sliding window was one SNP every 50 kb to ensure that low SNP density did not confound ROH detection. The maximum gap allowed between two consecutive homozygous SNPs in a ROH was 100 kb. Subsequently, the proportion of individuals with ROH was calculated for each sliding window. Regions enriched for ROH were defined as regions with top 1% of the empirical distribution of the frequency of a SNP occurring in ROH across individuals.

The EHH-based test statistics, including EHH surrounding the lead SNPs of detected QTL and |iHS| for each SNP, were calculated according to Sabeti et al. [22] and Voight et al. [23] using R package rehh [24, 25]. Ancestral and derived alleles were obtained for 539,043 SNPs of Illumina BovineHD Bead chip from Rocha et al. [26], where the ancestral allele was determined as the most frequent allele observed in yak, buffalo and sheep with a minimum of two copies in common across all species to avoid misclassifications due to sequencing errors. Phased haplotypes were obtained using *Beagle* (version 3.3.2) with 20 iterations of phasing algorithm [27]. The iHS values were standardized within derived allele frequencies bins of 0.02 according to Voight et al. [23]. The *P* value of the iHS score was derived as *P* = 1 − 2|Φ(iHS) − 0.5|, where Φ(iHS) is the cumulative Gaussian distribution function of iHS (under neutrality), and *P* is the two-tailed *P* value associated with the null hypothesis assuming no selection [28]. The QTL detected in the 99-trait meta-analysis were considered to be under selection if the respective lead SNPs were located in regions containing SNPs with *P* < 0.001.

## Results

We considered 4,578 Brown Swiss bulls for our analyses that had estimated breeding values (EBVs) for 56 economically important traits. The EBVs originated from genetic evaluations that were performed separately for the Austrian-German and Swiss populations. Analyzing phenotype data from the two populations jointly was not readily possible because the respective EBVs relate to different base populations and thus are not comparable. Moreover, traits may be defined slightly different in both populations. Thus, we considered the Austrian-German and Swiss populations as two separate cohorts and performed a total of 99 single-trait GWAS (44 traits in Austrian-German and 55 traits in Swiss population, see Additional file 1: Table S1 for more information about the traits considered). These 99 GWAS revealed a total of 1,067 significant (*P* < 8.36 x 10^−8^) trait x SNP associations. Of these, 648 and 305 SNPs were significantly associated with one and two traits, respectively, and 114 SNPs were significantly associated with at least three traits (Additional file 2: Fig. S1), suggesting that pleiotropic effects are present and detectable in the studied populations.

To increase the power of the association tests and characterize pleiotropic QTL, we carried out multi-trait meta-analyses with traits classified into eleven trait categories (referred to as trait-cat meta-analyses). Categories of traits were defined according to the classification of the traits at the Interbull Centre (http://www.interbull.org/ib/geforms), including milk production, body conformation (body size, leg confirmation and mammary gland morphology), reproduction (fertility and calving), and growth and carcass quality. The trait-cat meta-analyses also enabled us to combine similar traits from the two distinct cohorts, thus possibly increasing the power to detect QTL for traits with similar definitions, such as milking speed, longevity, persistence and somatic cell score. The number of single-trait analyses that were combined to meta-analyses ranged from two (for milking speed, longevity, persistence and somatic cell score) to 26 (for mammary gland morphology; see Additional file 1: Table S1 for the trait classifications).

### Summary of QTL effects and candidate genes by different categories of traits

The eleven trait-cat meta-analyses revealed a total of 1,203 SNPs that were significantly associated in at least one multi-trait meta-analysis, thus increasing by 13% the number of significantly associated SNPs compared to the single-trait GWAS. By grouping significant SNPs located within 5 Mb of the lead SNP, we identified between zero (somatic cell score and lactation persistency) and seven QTL (milk production) for the trait-cat meta-analyses. These QTL are characterized in more detail below separately for the trait categories considered (Fig. 1; Table 1).

**Figure 1.**
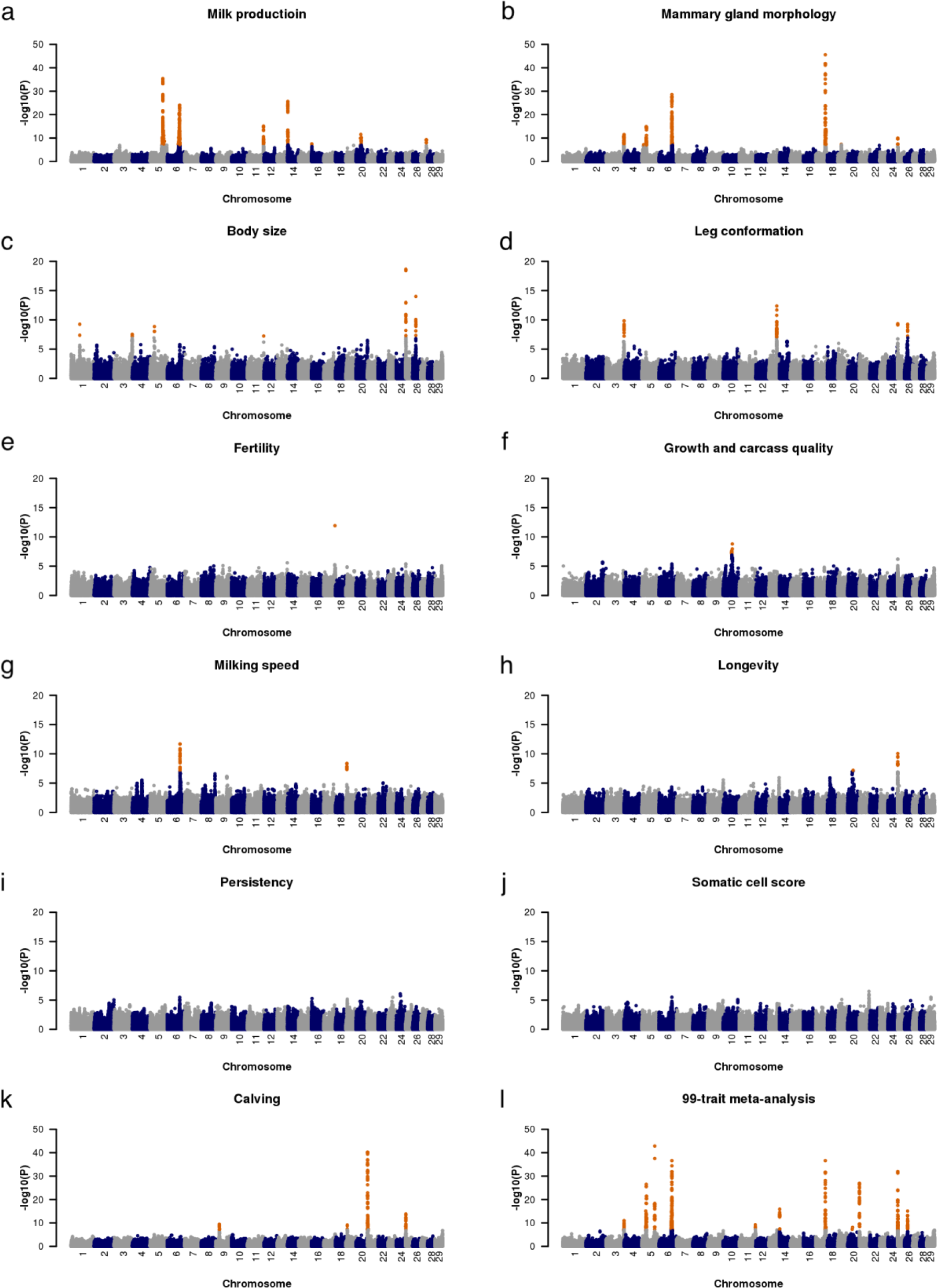
Manhattan plots for 11 trait-cat meta-analyses and 99-trait meta-analysis. Manhattan plot representing the association of 598,016 autosomal SNPs. Orange color represents variants with a P_META_ less than 8.36 x 10^−8^. The y-axis for the milk production meta-analysis was cut off at -log10(*P*) =50.

**Table 1.**
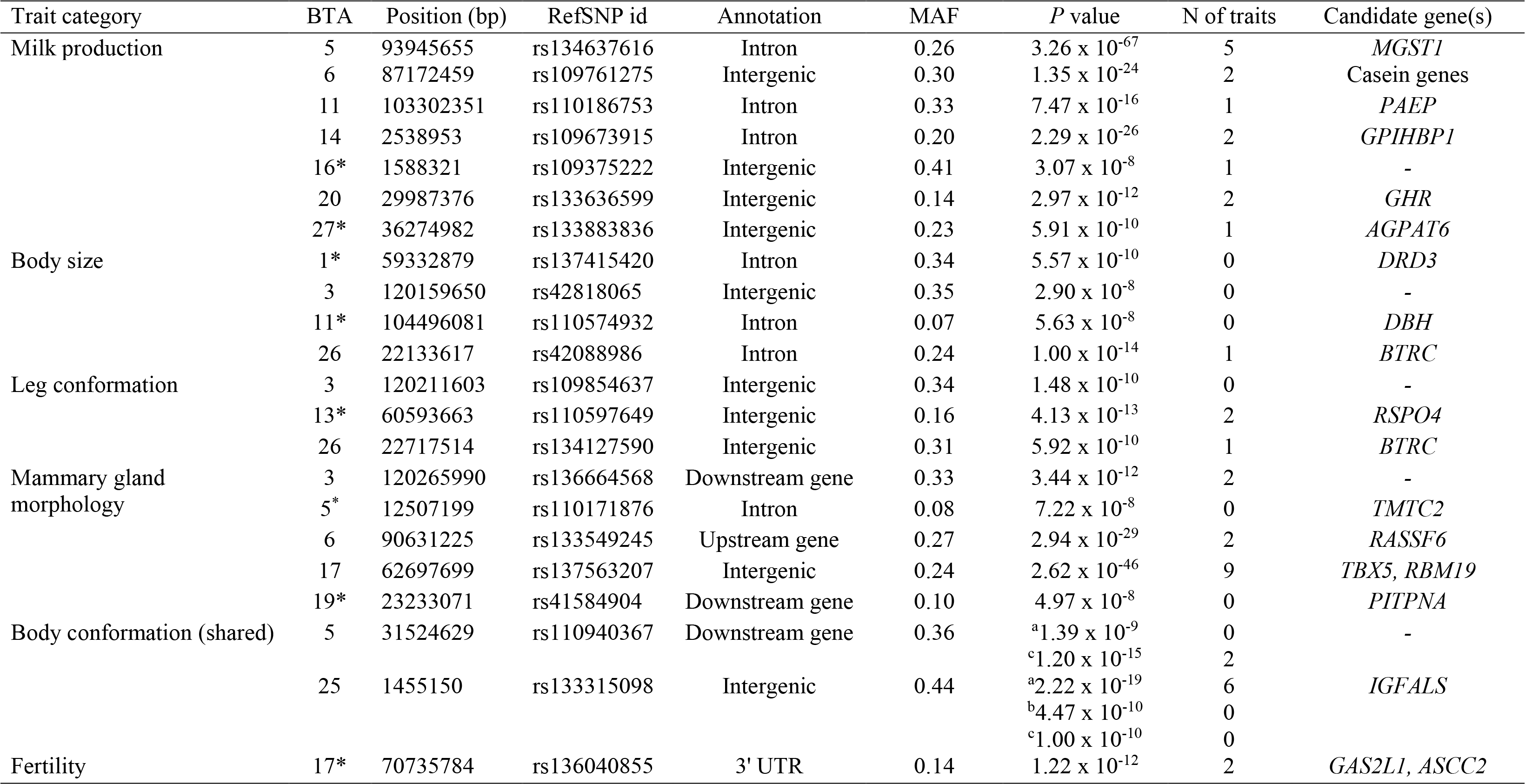
The most significant SNP at QTL detected across trait-cat meta-analyses and potential candidate genes.

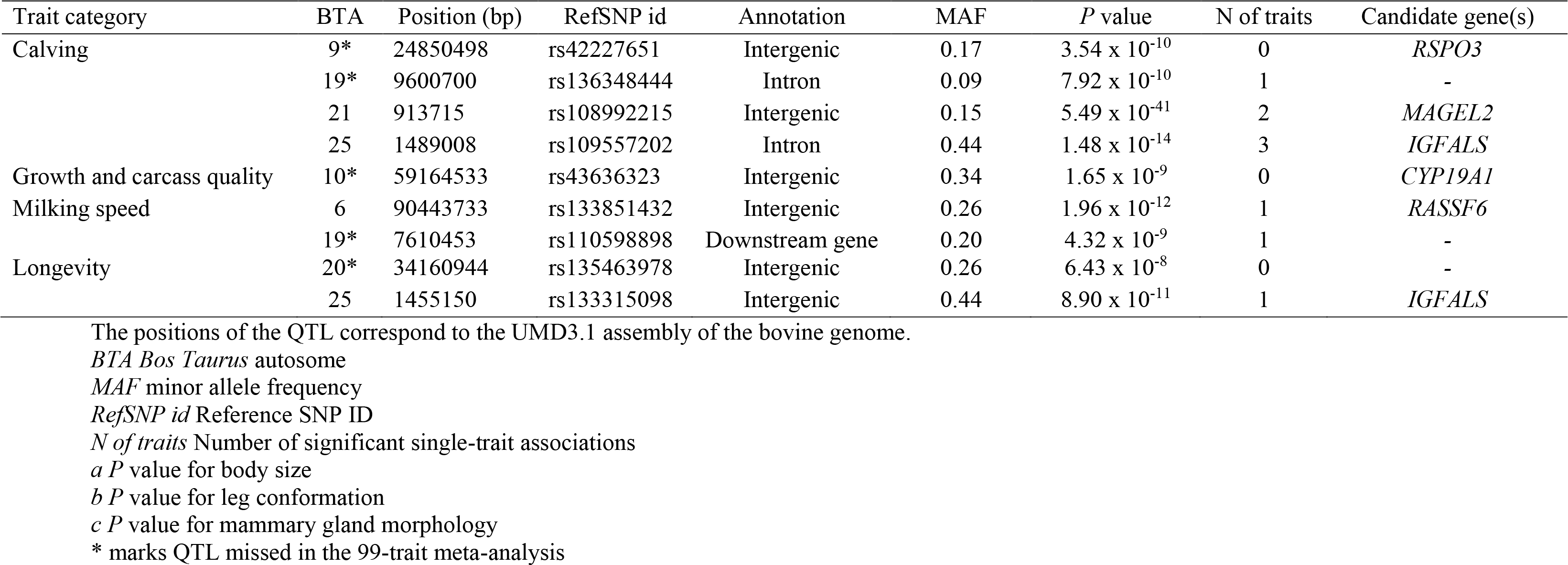

#### Milk production

We detected seven QTL (*P* < 8.36 x 10^−8^) that were associated with milk production traits on seven different chromosomes in Brown Swiss cattle. Of these, four QTL were significantly associated with multiple traits in the single-trait GWAS: a QTL on BTA5 was associated with five traits, and three QTL on BTA6, 14 and 20 were associated with two traits (Table 1). Five of seven milk production QTL encompass genes that had previously been reported to underpin milk yield or content in cattle, including the *MGST1* gene on BTA5 [11, 29], the cluster of casein genes on BTA6 [30], the *PAEP* (also known as β-lactoglobulin) gene on BTA11, the *GHR* gene on BTA20 [31] and the *AGPAT6* gene on BTA27 [32]. Note that the gene next to the lead SNP (rs133636599, at 29,987,376 bp) on BTA20 was not *GHR* but *MRPS30* encoding mitochondrial ribosomal protein S30, which we did not consider as an obvious candidate gene underpinning variation in milk production traits. The *GHR* gene encoding the growth hormone receptor was considered as a plausible candidate gene for the QTL on BTA20 because significantly associated SNPs were detected between 29.9 and 34.8 Mb, i.e., a region that coincided with a previously reported QTL for milk production traits in Brown Swiss cattle, where the *GHR* p.F279Y-variant [31] was the most significantly associated variant [11].

Another two QTL for milk production traits were located on BTA14 and BTA16. Both regions have recently been described to harbor QTL for milk production traits in several cattle breeds including Brown Swiss [11, 33, 34]. The well-known *DGAT1* p.A232K-variant [35] can be excluded as the causal variant for the milk production QTL on BTA14 because it does not segregate in Brown Swiss cattle [11]. For a QTL for milk production traits in close vicinity to *DGAT1* on BTA14, the *GPIHBP1* gene has been postulated as candidate gene [34, 36].

#### Body conformation

A total of 60 traits related to body conformation (30 traits for each cohort) were classified into three categories, i.e., body size, leg conformation and mammary gland morphology. The respective three trait-cat meta analyses revealed six, four and seven QTL that were associated with body size, leg conformation and mammary gland morphology, respectively (Fig.1; Table 1). Of six QTL associated with body size, four were not significantly associated with any of the 99 traits in the single-trait GWAS, i.e. QTL on BTA1, 3, 5 and 11. A QTL on BTA25 was associated with six traits related to body size, and a QTL on BTA26 was associated with one trait related to body size in the single-trait GWAS. Of four QTL associated with leg conformation, two were not detected in the single-trait GWAS, i.e. QTL on BTA3 and 25. Two QTL on BTA13 and 26 were associated with two and one trait(s) related to leg conformation, respectively. Of seven QTL associated with mammary gland morphology, three were not detected in single-trait GWAS, i.e. QTL at BTA5:12,507,199, 19 and 25. A QTL for mammary gland morphology on BTA17 was significantly associated with nine traits related to the shape of the mammary gland, and three QTL were associated with two traits.

Among the 17 QTL detected for three categories related to body conformation, four QTL on BTA3, 5, 25 and 26 were significant for more than one trait category. A QTL for body confirmation traits was located on BTA3. Significantly associated SNPs were detected in an interval between 117.3 and 120.2 Mb. This QTL was associated with body size (*P* = 2.90 x 10^−8^), leg confirmation (*P* = 1.48 x 10^−10^) and mammary gland morphology (*P* = 3.44 x 10^−12^). The lead SNPs differed for the three trait categories but were in very high linkage disequilibrium (LD; 0.93 < r^2^ < 0.97). This region coincides with a QTL for stature that has been detected in various cattle breeds [37]. A total of 12 annotated genes were located within a 1-Mb interval centered on the lead SNPs at this QTL. However, we did not observe an obvious candidate gene that might underpin variation in body conformation traits.

Another QTL for body confirmation traits was located on BTA5 with significantly associated SNPs being located between 29.9 and 31.5 Mb. This QTL was associated with body size (*P* = 1.39 x 10^−9^) and mammary gland morphology (*P* = 1.20 x 10^−15^), and the same SNP was the lead SNP (rs110940367 at 31,524,629 bp) for both trait categories. A total of 25 genes were annotated within 1 Mb of the lead SNP, including 11 protein-coding genes. However, none of the genes had obvious functions related to growth or mammary gland development.

A QTL for body conformation was located at the proximal region of BTA25, and the same SNP (rs133315098 at 1,455,150 bp) was the lead SNP for all three trait categories related to body conformation (*P* = 2.22 x 10^−19^ for body size; *P* = 4.47 x 10^−10^ for leg conformation; *P* = 1.00 x 10^−10^ for mammary gland morphology). This region had been previously shown to be associated with stature in Brown Swiss cattle [8]. A total of 68 annotated genes located within 1 Mb of the lead SNP, including 51 protein-coding genes. The closest gene to the lead SNP was *MEIOB* encoding the meiosis specific protein with OB function (19 KB upstream of the transcription start), which we did not consider as an obvious candidate gene for body conformation traits because this gene is mainly involved in meiotic recombination [38]. A potential candidate gene mapped to this region was *IGFALS* encoding insulin-like growth factor binding protein acid labile subunit, of which the lead SNP is located at about 87 KB upstream of the transcription start. Variation in the *IGFALS* gene has previously been shown to underpin growth-related traits in human and mouse [39].

Another QTL for body confirmation was located on BTA26 with significantly associated SNPs located between 21.6 and 22.7 Mb. It was associated with body size (*P* = 1.00 x 10^−14^) and leg conformation (*P* = 5.92 x 10^−10^). The lead SNPs differed between two trait categories but were in high LD (r^2^ = 0.70). So far, this region has not been reported to underpin body conformation traits in cattle. A total of 22 genes were annotated within 1 Mb of the lead SNPs, including 20 protein-coding genes. The lead SNP for body size (rs42088986 at 22,133,617 bp) was located within the third intron of *BTRC* encoding a member of the F‐box protein family. This gene is involved in Wnt signaling that plays a critical role in developmental processes and has been shown to be associated with limb development [40] and limb abnormalities [41].

Apart from QTL that were associated with several trait categories related to body conformation, we detected six trait-cat specific QTL, i.e., two QTL on BTA1 and 11 for body size, one QTL on BTA13 for leg conformation, and three QTL on BTA6, 17 and 19 for mammary gland morphology. These regions have previously been associated with various body conformation traits in several cattle breeds including Brown Swiss [9, 37, 42–44].

#### Reproduction (fertility and calving)

A QTL (*P* = 1.22 x 10^−12^) on BTA17 was associated with two fertility-related traits, i.e. interval from first to last insemination and non-return rate in heifers, and was also significant in the fertility meta-analysis. An association between genetic variants in this region and fertility has been reported previously in Brown Swiss cattle [10]. The lead SNP (rs136040855 at 70,735,784 bp) was located at the 3’UTR of the *AP1B1* gene encoding adaptor related protein complex 1 subunit beta 1, which we did not consider as a plausible candidate gene for fertility. A previous study revealed that two missense variants located in the *GAS2L1* gene encoding growth arrest specific 2 like 1 (at 70,724,328 bp) and *ASCC2* gene encoding activating signal cointegrator 1 complex subunit 2 (at 71,084,044 bp) segregate in Brown Swiss cattle, and the variant in *ASCC2* gene has been suggested as a plausible candidate causal mutation controlling female fertility in cattle [10].

Four QTL on BTA9 (*P* = 3.54 x 10^−10^), 19 (*P* = 7.92 x 10^−10^), 21 (*P* = 5.49 x 10^−41^) and 25 (*P* = 1.48 x 10^−14^) were associated in a meta-analysis of traits related to the calving performance. Three QTL on BTA19, 21 and 25 were significant in one, two and three single-trait GWAS, respectively. Variants nearby these regions have been shown to be associated with calving traits either in Brown Swiss or Fleckvieh cattle [10, 45, 46]. A QTL for calving traits on BTA9 was significant in the trait-cat meta-analysis but not in the single-trait GWAS. This region has not been reported so far to be associated with calving performance in cattle. Four genes were annotated within 1 Mb of the lead SNP (rs42227651 at 24,850,498 bp), including two protein-coding genes. The *RSPO3* gene, of which the lead SNP was located approximately 0.5 Mb downstream of the transcription end, might be considered as a potential candidate gene that might underpin variation in calving traits. This gene encodes R-spondin 3 which modulates WNT signaling [47] and is involved in blood vessel formation including placental development, which could affect fetal growth and thus result in calving difficulties [48].

#### Growth and carcass quality

The trait-cat meta-analysis revealed a QTL (*P* = 1.65 x 10^−9^) on BTA10 that was associated with growth traits and carcass quality. This QTL was not significant in the single-trait GWAS. The lead SNP (rs43636323) was located between the *GLDN* gene encoding gliomedin (about 33 KB upstream of the transcription start) and the *CYP19A1* gene encoding cytochrome P450, family 19, subfamily A, polypeptide 1 (about 63 KB upstream of the transcription start). A QTL for carcass traits and growth index in Holstein and Red Dairy cattle is located in immediate vicinity of the lead SNP and *CYP19A1* gene was considered as the candidate gene because it catalyzes the conversion of androgens to estrogens [49]. Variation in the *CYP19A1* gene is associated with both growth and reproduction in mice and humans [50, 51].

#### Management (Milking speed and longevity)

We detected two QTL on BTA6 (*P* = 1.96 x 10^−12^) and 19 (*P* = 4.32 x 10^−9^) for milking speed and two QTL on BTA20 (*P* = 6.43 x 10^−8^) and 25 (*P* = 8.90 x 10^−11^) for longevity (Fig.1; Table 1). The QTL on BTA6 and 25 were also associated with body conformation traits (described above). The lead SNP of a longevity QTL on BTA25 was also the lead SNP for body conformation traits. A QTL for milking speed on BTA19 coincides with a QTL for milking speed in Nordic Holstein [52]. Ten genes were annotated within 1 Mb of the lead SNP (rs110598898 at 7,610,453 bp), including seven protein-coding genes. However, we did not detect an obvious candidate gene related to milking speed. A QTL for longevity was located on BTA20 at about 34 Mb. This QTL was not detected in the single-trait GWAS. However, we did not observe an obvious candidate gene that might underpin variation in longevity.

#### Overview of QTL segregating in Brown Swiss cattle

In an attempt to compile an overview of major QTL segregating in Brown Swiss cattle, we performed a multi-trait meta-analysis combining the results of all 99 single-trait GWAS. The 99-trait meta-analysis revealed 531 significantly associated SNPs that clustered at 12 QTL (*P* < 8.36 x 10^−8^) regions on BTA3, 5, 6, 11, 14, 17, 20, 21, 25 and 26. When compared to the trait-category based meta-analyses, the 99-trait meta-analysis did not reveal any new QTL (Fig. 1) and less than half the number of SNPs was significantly associated. Moreover, 13 QTL that were detected in at least one trait-cat meta-analysis were not significantly associated in the 99-trait meta-analysis (Table 1). The maximum proportion of variance in daughter-derived phenotypes explained by the QTL detected in 99-trait meta-analysis across traits ranged from 0.02 for the QTL on BTA11 to 0.10 for the QTL at BTA5:93,945,655 both for fat percentage (Table 2).

**Table 2.** The properties of 12 QTL detected in the 99-trait meta-analysis.

| BTA | Position (bp) | RefSNP id | Annotation | MAF | Effect allele | Ancestral allele | *P* value | *V<sub>p</sub>* | iHS | Trait category | Candidate gene(s) |
| --- | --- | --- | --- | --- | --- | --- | --- | --- | --- | --- | --- |
| 3 | 120211603 | rs109854637 | Intergenic | 0.34 | A | A | $1.02 \times 10^{-11}$ | 0.022 | - | body conformation | - |
| 5 | 31524629 | rs110940367 | Downstream gene | 0.36 | G | G | $3.18 \times 10^{-27}$ | 0.020 | -1.65 | body conformation | - |
| 5 | 93945655 | rs134637616 | Intron | 0.26 | C | C | $1.35 \times 10^{-43}$ | 0.100 | 1.35 | milk production | *MGST1* |
| 6 | 87005244 | rs135679764 | Intron | 0.18 | G | - | $1.87 \times 10^{-15}$ | 0.032 | - | milk production | Casein genes |
| 6 | 90486780 | rs41654962 | Intergenic | 0.20 | G | G | $2.35 \times 10^{-37}$ | 0.071 | 2.49 | body conformation, milking speed | *RASSF6* |
| 11 | 103302351 | rs110186753 | Intron | 0.33 | G | - | $6.64 \times 10^{-10}$ | 0.020 | - | milk production | *PAEP* |
| 14 | 2534899 | rs109673915 | Intron | 0.25 | G | G | $1.33 \times 10^{-16}$ | 0.046 | - | milk production | *GPIHBP1* |
| 17 | 62697699 | rs137563207 | Intergenic | 0.24 | C | C | $2.35 \times 10^{-37}$ | 0.035 | 0.19 | body conformation | *TBX5, RBM19* |
| 20 | 29987376 | rs133636599 | Intergenic | 0.14 | A | C | $8.24 \times 10^{-9}$ | 0.027 | 1.21 | milk production | *GHR* |
| 21 | 913715 | rs108992215 | Intergenic | 0.15 | A | - | $1.30 \times 10^{-27}$ | 0.036 | - | calving | *MAGEL2* |
| 25 | 1455150 | rs133315098 | Intergenic | 0.44 | A | A | $9.72 \times 10^{-33}$ | 0.032 | - | body conformation, calving, longevity | *IGFALS* |
| 26 | 22133617 | rs42088986 | Intron | 0.24 | A | A | $8.59 \times 10^{-16}$ | 0.025 | -0.57 | body conformation | *BTRC* |

The positions of the QTL correspond to the UMD3.1 assembly of the bovine genome.
*BTA* *Bos Taurus* autosome
*MAF* minor allele frequency
*RefSNP id* Reference SNP ID
*V<sub>p</sub>* maximum proportion of variance in daughter-derived phenotypes explained by the QTL across traits
*Trait type* Trait group with which the QTL were associated
*iHS* integrated haplotype score (not available for SNPs located at the beginning and the end of the chromosome)
- not available

Of these 12 detected QTL, five, six and one were associated with milk production, body conformation and calving performance, respectively (Table 2). The 99-trait meta-analysis enabled us to distinguish between two neighboring QTL on BTA6: a QTL at BTA6:87,005,244 associated with milk production traits and a QTL at BTA6: 90,486,780 associated with mammary gland morphology and milking speed (Additional file 3: Fig. S2). Compared to the trait-cat meta-analyses, the lead SNPs of the 99-trait meta-analysis differed for two QTL on BTA6 and one QTL on BTA14. The lead SNP at BTA6:87005244 in the 99-trait meta-analysis was about 0.17 MB apart from the one for milk production (at 87,172,459 bp), but they were in moderate LD (r^2^ = 0.41). The lead SNP at BTA6:9048678 in the 99-trait meta-analysis was located between the lead SNPs for milking speed (at 90,443,733 bp) and mammary gland morphology (at 90,631,225 bp). These three SNPs were in moderate LD (0.51 < r^2^ < 0.70). The lead SNP at a QTL on BTA14 (at 2,534,899 bp) in the 99-trait meta-analysis was about 4 Kb distant from the lead SNP of the milk production (at 2,538,953 bp), but they were in high LD (r^2^ = 0.75). Furthermore, four QTL that were detected in more than one trait-cat meta-analysis, i.e. QTL at BTA5:31524629, BTA6:90,486,780, BTA25:1,455,150 and BTA26:22,133,617, had lower *P* values in the 99-trait meta-analysis than any of the trait-cat meta-analyses, whereas all other QTL that were significant for only one trait category had higher *P* value in the 99-trait meta-analysis than any of the trait-cat meta-analyses (Table 1 and 2).

Two QTL were shared across different trait categories that were not related to body conformation, i.e. QTL at BTA6:90,486,780 and BTA25:1,455,150 (Table 2). The QTL on BTA6 was associated with mammary gland morphology and milking speed. The *G* allele of the lead SNP (rs41654962) had a frequency of 0.8 in Brown Swiss cattle. It was associated with thinner teats and reduced milking speed (Table 2; Additional file 4: Table S2). The QTL on BTA25 was associated with body size, leg confirmation, mammary gland morphology, calving ease and longevity. The *A* allele of the lead SNP (rs133315098) had a frequency of 0.56 and was associated with high birth weight (both direct and maternal), greater stature and longer gestation length. It was also associated with calving difficulties (both direct and maternal), greater stillbirth incidence and reduced longevity (Table 2; Additional file 5: Fig. S3).

### Selection signatures

We explored if recent artificial selection for economically important traits left detectable traces in the Brown Swiss cattle genome. To detect evidence of selection, we applied two test statistics: runs of homozygosity (ROH) and the integrated haplotype score (|iHS|). We also calculated expected haplotype homozygosity (EHH) surrounding the lead SNPs of QTL detected from the 99-trait meta-analysis as well as assessed if their allele frequencies changed over time, thus investigating if these QTL were targets of recent or ongoing selection.

#### Genome-wide scans of selection signatures

Regions on BTA4, 5, 6, 11, 16, 19 harbored ROH that were frequent in Brown Swiss cattle. However, the lead SNPs of QTL detected from 99-trait meta-analysis were rarely located in ROH that were shared across individuals except for the ones on BTA6 and 25 (Fig. 2).

**Figure 2.**
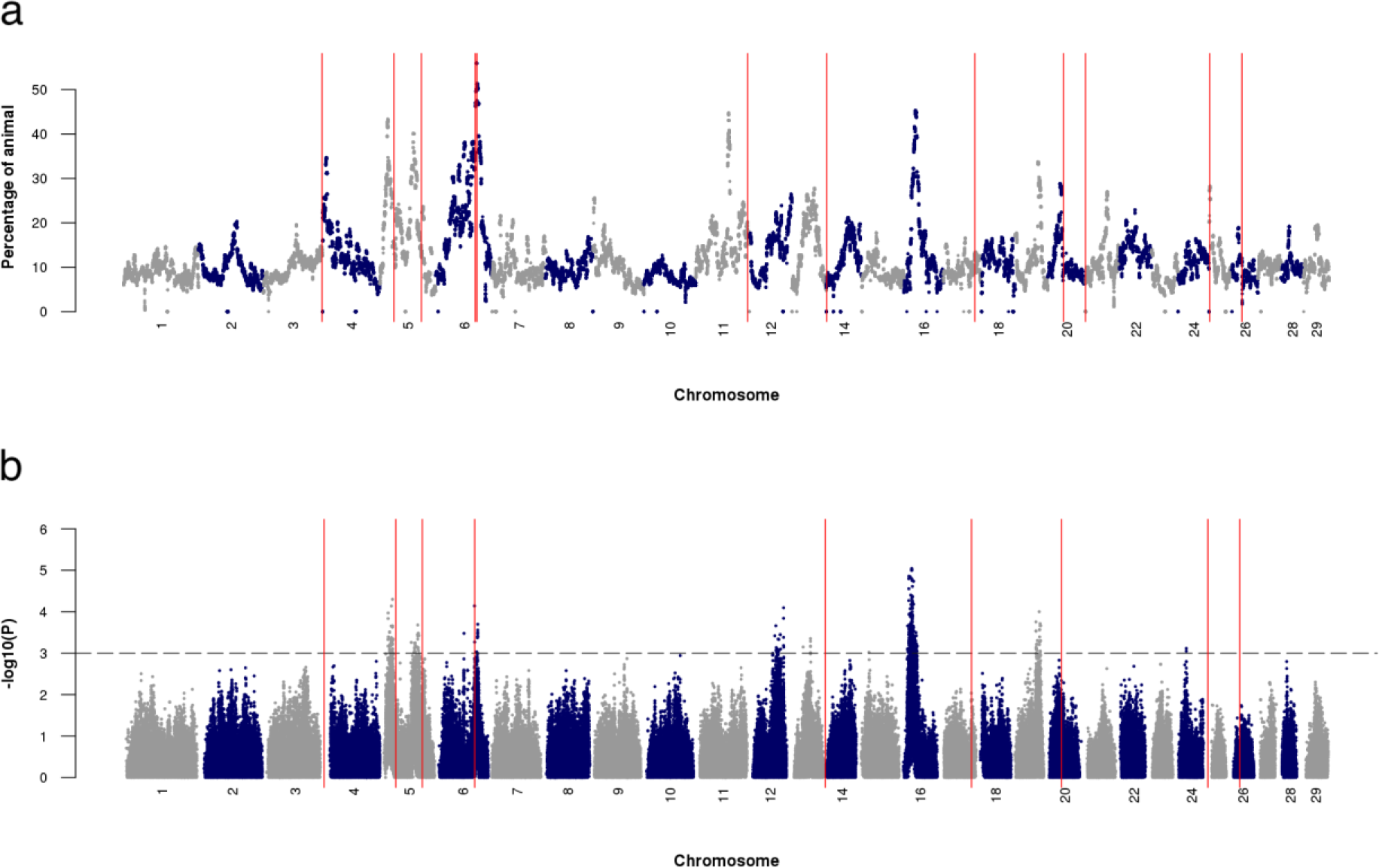
Selection signatures detected by runs of homozygosity (a) and integrated haplotype score (b). Red vertical lines mark the locations of the lead SNPs at QTL detected from the 99-trait meta-analysis. The horizontal line represents the genome-wide significant level of integrated haplotype score test (*P* < 0.001). Note that the integrated haplotype scores were not available for SNPs located at the beginning and the end of the chromosome to prevent edge effects.

We calculated iHS scores for each SNP to detect ongoing selection at alleles that have not yet reached fixation. Negative and positive iHS values indicate selection for derived and ancestral alleles, respectively. Evidence of strong selection (*P* < 0.001) was found for genomic regions on BTA5, 6, 12, 13, 14, 15, 16, 19 and 24 (Fig. 2), including five regions on BTA5, 6, 12, 16 and 19 that were previously detected in a smaller sample of Brown Swiss cattle [53]. The lead SNPs of QTL detected in the 99-trait meta-analysis did not coincide with candidate regions of selection signature except for the lead SNP at BTA6:90,486,780. Ten SNPs on BTA6 (located between 90.2 and 91.7 Mb) were significant both in the 99-trait meta-analysis and iHS test. Note that iHS values were not calculated for SNPs located either at the proximal or distal regions of chromosomes to prevent edge effects [24].

#### Signatures of selection at the lead SNPs of the multi-trait meta-analysis

Derived and ancestral alleles could be determined for the lead SNPs at nine out of twelve QTL detected in the 99-trait meta-analysis. Generally, all lead SNPs showed either no or only weak evidence of selection because most of their corresponding |iHS| values were less than 1.98 (Table 2). However, the |iHS| value was 2.49 for the lead SNP at BTA6:90,486,780 that was associated with mammary gland morphology and milking speed, suggesting that this QTL is a target of ongoing selection. A positive iHS value at the lead SNP indicates selection for the ancestral allele that was associated with thinner teats and reduced milking speed (Additional file 4: Table S2). This is also shown by a slower EHH decay over the distance surrounding the ancestral allele than the derived allele of this SNP (Fig. 3).

**Figure 3.**
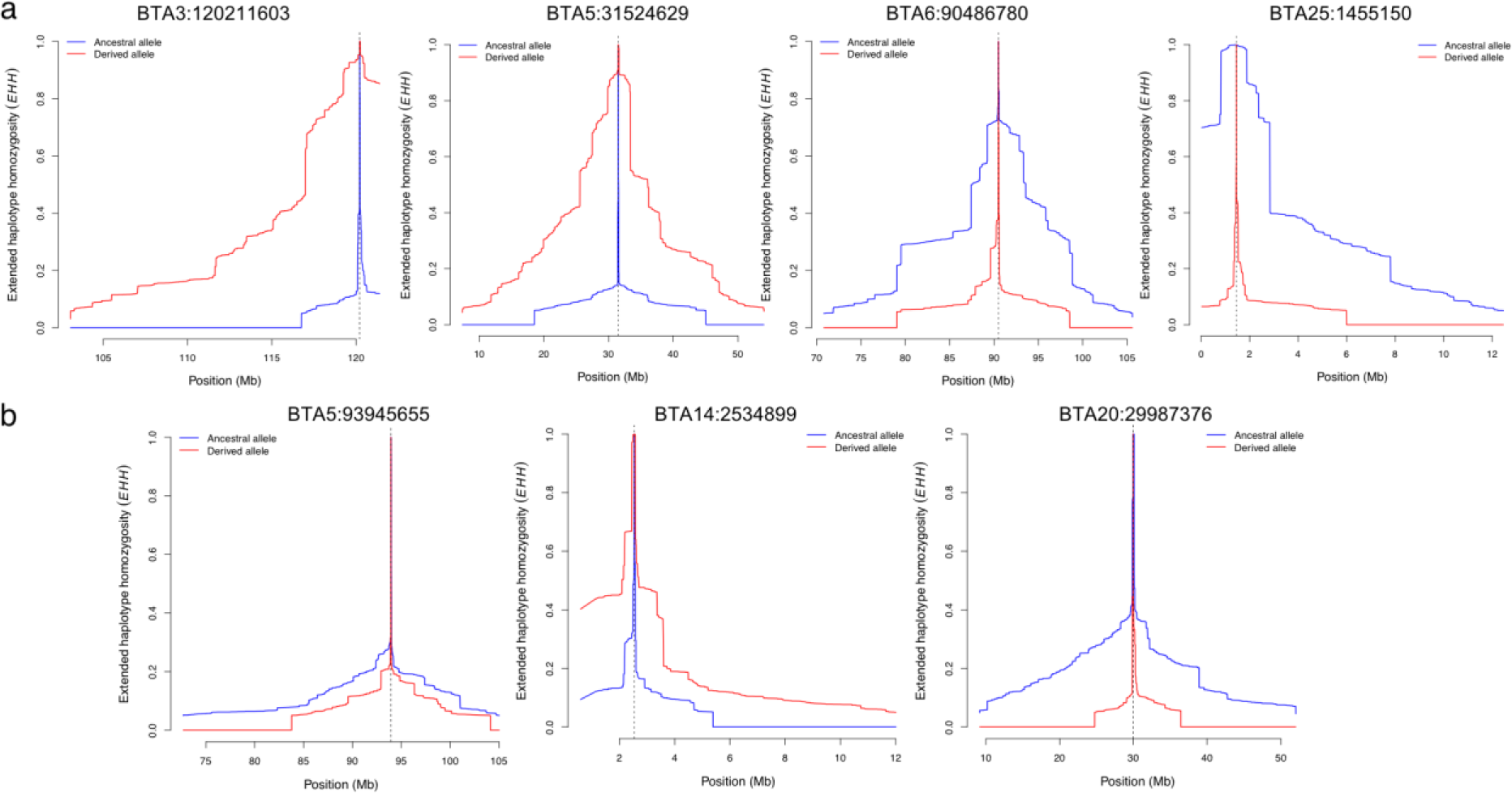
Extended haplotype homozygosity (EHH) decay surrounding the lead SNPs at QTL affecting body conformation and milk production. a. the lead SNPs of the QTL associated with body conformation b. the lead SNPs of the QTL associated with milk production, where the ancestral alleles could be determined. The dashed line marks the location of the lead SNPs relative to which EHH scores were calculated. Blue and red lines indicate EHH decay of the ancestral and derived alleles, respectively.

Of three lead SNPs (BTA5:31,524,629, BTA5:93,945,655 and BTA20:29,987,376) that showed weak evidence of selection (|iHS| > 1), the lead SNP at BTA5:31,524,629 that was associated with body size and mammary gland morphology showed a markedly different decay of haplotype homozygosity surrounding the ancestral and derived alleles (Fig. 3). Among three lead SNPs (BTA3:120,211,603, BTA14:2,534,899 and BTA25:1,455,150) that were located in proximal and distal regions of chromosomes, the lead SNP at BTA3:120,211,603 and BTA25:1,455,150 that were associated with body conformation also showed difference in decay of haplotype homozygosity surrounding the ancestral and derived alleles compared to the lead SNP at BTA14:2,534,899 that was associated with milk production (Fig. 3). The alleles at BTA3:120,211,603, BTA5:31,524,629 and BTA25:1,455,150 that increased body size showed slower EHH decay over distance than the alleles with the opposite effects (Fig. 3; Additional file 4: Table S2).

Six out of twelve lead SNPs detected in 99-trait meta-analysis showed significant changes (*P*_*Bonferroni-corrected*_ < 0.05) in their allele frequencies from birth year 1990 to 2015 (Fig. 4). Out of these six lead SNPs, four were associated with body conformation traits. The alleles that increased in frequency over time were associated either with larger body size or thinner teats (Additional file 4: Table S2). The other two lead SNPs were associated with milk production, and the alleles that increased in frequency over time were associated with higher protein and fat percentage (Additional file 4: Table S2).

**Figure 4.**
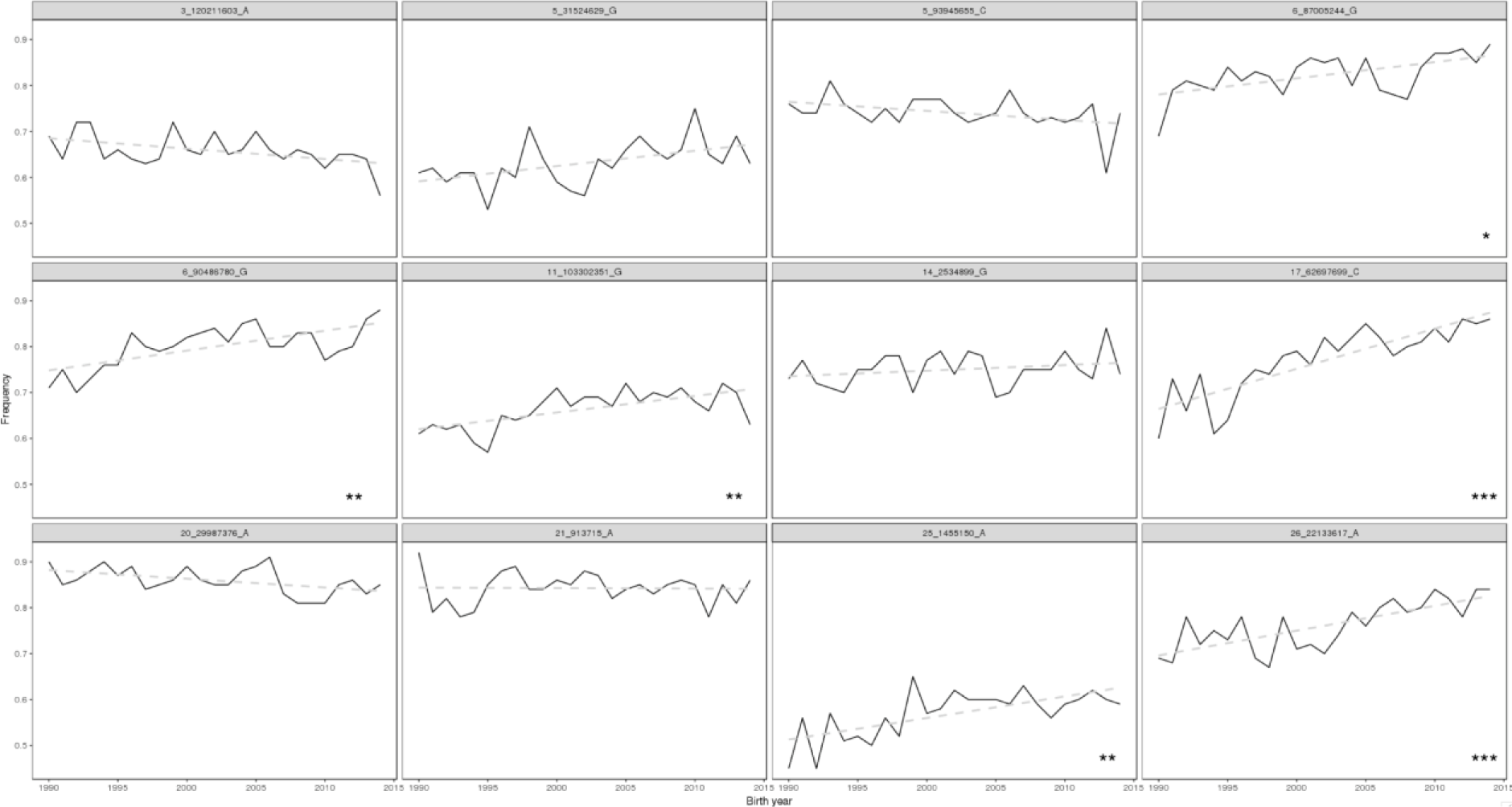
Changes in effect allele frequencies of the lead SNPs at QTL detected from 99-trait meta-analysis. *,**,*** mark the significant level of changes in allele frequency (*P*_*Bonferroni-corrected*_ < 0.05, *P*_*Bonferroni-corrected*_ < 0.01, *P*_*Bonferroni-corrected*_ < 0.001).

## Discussion

We performed single-trait GWAS between partly imputed genotypes at 598,016 SNPs and 99 daughter-derived phenotypes for economically important traits in 4,578 Brown Swiss bulls from Austria, Germany and Switzerland. Subsequently, we performed multi-trait meta-analyses of eleven trait categories and a multi-trait meta-analysis that combined all 99 traits to provide a global overview of major QTL segregating in European Brown Swiss cattle. In contrast to Guo et al. [8], we did not analyze the daughter-derived phenotypes from these two populations jointly because we used records that were obtained from distinct national genetic evaluations that differ in trait definitions, scales and base populations. Using multi-trait across country evaluations (MACE) on the national records would provide comparable daughter-derived phenotypes for bulls from different populations [54, 55]. However, since MACE were not available for the Brown Swiss bulls of our study, we applied a step-wise procedure to consider all animals in our dataset for the association analyses: we first carried out single-trait GWAS for the two populations separately and subsequently combined similar traits in an across-cohort multi-trait meta-analysis. The number of significantly associated SNPs was clearly higher in the trait-category-specific meta-analyses than single-trait GWAS, indicating that this strategy increased the power to detect QTL. Our results corroborate previous findings that multi-trait meta-analysis provides high power to detect QTL, particularly for correlated traits [3, 44, 56]. However, when we combined all 99 single-trait GWAS using a multi-trait meta-analysis, the number of significantly associated SNPs and detected QTL was lower than in the trait-cat meta-analyses. Our results, obtained from a moderately sized mapping population, indicate that the 99-trait meta-analysis had less power than the trait-cat meta-analyses to detect QTL that affect only one trait category. This finding is supported by increased *P* values for seven out of twelve QTL that were significant in 99-trait meta-analysis but affect only one trait category. Therefore, our results suggest that appropriate clustering of traits with similar definitions or pronounced genetic correlations may provide higher power to detect QTL than meta-analysis of a larger number of (partly) uncorrelated traits.

We detected 25 major QTL (*P* < 8.36 x 10^−8^) affecting economically important traits using 4,578 Brown Swiss bulls that had both daughter-derived phenotypes and partially imputed genotypes at 598,016 autosomal SNPs. In spite of a decent sample size and large number of traits with relatively high heritability [57], the number of detected QTL was rather low in our study possibly reflecting a highly polygenic nature of most of the traits considered. It seems likely that most of the economically important traits in Brown Swiss cattle are predominantly determined by many small-effect loci that remained undetected in our study. Our study revealed QTL that explained between 2 and 10% of the trait variation. Increasing the marker density, e.g., by imputing sequence variant genotypes might facilitate the fine mapping of the detected QTL [33] and uncover additional QTL that have not been detected using microarray-derived genotypes [58]. However, the markers on the BovineHD Bead chip are likely dense enough to detect QTL in the most popular breeds including Brown Swiss cattle [59]. Substantially increasing the number of genotyped animals seems to be required to detect QTL that explain a smaller proportion of the trait variation in the Brown Swiss cattle. Considering that the population size is considerably smaller in Brown Swiss than Holstein cattle, either applying MACE on progeny-tested animals from distinct populations or including cows with own performance records will increase the size of the mapping populations and thus the power to detect QTL. Moreover, estimating the effects of all markers simultaneously might facilitate to detect QTL with smaller effects [60].

Most of the QTL detected in our study were located nearby QTL that had been described previously in several cattle breeds including Brown Swiss, thus confirming the relevance of these genomic regions for shaping economically important traits in cattle. Most QTL in the 99-trait meta-analysis are associated with milk production, body conformation and, to a lesser extent, calving performance, possibly reflecting that the heritability and thus power to detect associations is higher for production than functional traits. In agreement with findings in Holstein cattle, most of the pleiotropic QTL detected in our study in Brown Swiss cattle were related to body conformation traits [42, 43], likely indicating common physiological pathways controlling different body features across cattle breeds. We did not detect pleiotropic QTL that were associated with apparently uncorrelated trait categories, such as milking speed and fertility possibly due to lack of power and stringent significant threshold (Bonferroni correction).

Among 25 detected QTL, five were significantly associated with at least two distinct traits across populations, i.e. QTL at BTA5:93,945,655, BTA6:90,486,780, BTA17:62,697,699, BTA21:913,715 and BTA25:1,455,150, indicating widespread pleiotropy. A larger mapping population might uncover pleiotropic effects at more QTL. Of these five QTL, QTL at BTA5:93,945,655, BTA17:62,697,699 and BTA21:913,715 affect several traits in the same trait category, whereas QTL at BTA6:90,486,780 and BTA25:1,455,150 affect a broad range of traits from different categories. A QTL at BTA5 encompassing the *MGST1* gene affects fat percentage, fat yield and protein percentage. Its pleiotropic effects on milk production traits have been shown for several breeds [11, 29]. The QTL at BTA17:62,697,699 affects rear udder width, fore udder length, front and rear teat placement, and udder cleanness. Its effects on mammary gland morphology have been shown previously in Brown Swiss [9] and Fleckvieh cattle [44].

For two QTL (at BTA6:90,486,780 and BTA25:1,455,150) affecting a broad range of economically important traits, we observed lower *P* values of the lead SNPs from the 99-trait meta-analysis compared to trait-cat meta-analyses. Such a pattern could indicate that they underpin variation in traits from different categories. The QTL at BTA6:90,486,780 affects mammary gland morphology and milking speed. Our results show that this QTL has a larger effect on teat thickness than any other morphological feature of the mammary gland (see Additional file 4: Table S2). This genomic region has previously been identified as a major QTL for mammary gland morphology in Brown Swiss [9], Fleckvieh [44] and Holstein [61] cattle. Our results show that an allele of the lead SNP is associated with thinner teats (*P* = 2.94 x 10^−29^) and reduced milking speed (*P* = 1.96 x 10^−12^), indicating pleiotropic effects that had not been detected so far. It seems plausible that the anatomy of the teat could affect the milking speed, i.e., milk flow is faster in cows with wider teat canal and sphincter [62], which is also shown by the positive correlation between the teat thickness and milking speed [63]. An ideal shape of the mammary gland and an average milking speed are important for efficient machine milking and healthy udders [62, 64, 65]. This implies that bulls with EBVs around the population average are required for the breeding of cows with optimum teat thickness and milking speed. We observed that the cows’ teats became thinner within the past 25 years as a part of improving mammary gland morphology in Brown Swiss cattle while the milking speed increased considerably (see Additional file 6: Fig S4 and [66]). The increase in milking speed may result from accumulated positive effects of small-effect QTL that had not been detected in our study. Additionally, given that the QTL at BTA6:90,486,780 explains considerably more variation in teat thickness (5% for Swiss and 7% for Austrian-German population) than milking speed (2% for Swiss and 0.1% for Austrian-German population), selecting for the QTL at BTA6:90,486,780 would affect teat thickness to a larger extent than milking speed.

A QTL at BTA25:1,455,150 affects birth weight, body size, gestation length, calving ease and longevity. This QTL has been reported for stature and calving performance in Brown Swiss cattle [8, 10]. Our results show that this QTL also underpins variation in longevity and thus might contribute to the overall genetic correlations among stature, calving performance and length of productive life. As shown by the patterns of QTL effects across traits, the allele that is associated with larger body size and calving difficulties is also associated with longer gestation length. It is likely that this QTL acts on both pre- and postnatal growth. Enhanced fetal growth may cause dystocia and higher stillbirth incidence, and adult size affects calving abilities of cows [67, 68]. Since the QTL affects both pre- and postnatal growth, selection for an increased adult size and/or birth weight might lead to greater calving difficulties and subsequently higher calf mortality and reduced length of productive life of cows. Nonetheless, traits that are unfavorably correlated with calving performance (i.e. pre- and postnatal growth) could respond favorably by applying appropriate weights in a selection index or multi-trait BLUP [69, 70].

Only one QTL at BTA6:90,486,780 showed evidence of ongoing selection (based on ROH and iHS statistics). Consistent with previous findings [19, 71], we did not detect compelling evidence for the presence of selection signatures around the detected QTL although strong artificial selection particularly on milk production and body conformation traits took place recently in Brown Swiss cattle [66]. Each of the QTL detected in our study explains only a small fraction (less than 5% except for the one at BTA5:93,945,655) of variance in daughter-derived phenotypes for a given quantitative trait. Moreover, the QTL effects are likely overestimated due to Winner’s curse [72], indicating that the proportion of variance in daughter-derived phenotypes explained by a QTL is less than estimated in this study. Therefore, ongoing selection for economically important traits acts on a multitude of genomic regions and is unlikely to change the allele frequency of individual QTL rapidly as shown by the little changes in allele frequencies for the most of lead SNPs at QTL affecting milk production over the past 25 years. Nevertheless, QTL associated with body conformation, i.e. QTL at BTA3:120,211,603, BTA5:31,524,629 and BTA25:1,455,150 showed increased homozygosity surrounding the favorable alleles compared to the lead SNPs associated with milk production, confirming recent selection for physical appearance and mammary gland morphology in Brown Swiss cattle, especially for increasing body size and decreasing teat thickness [37, 66, 73]. This is further supported by the observed significant changes in frequency of the favorable alleles of the lead SNPs at four out of six QTL associated with body conformation.

## Conclusions

Here we report an overview of QTL affecting economically important traits in Brown Swiss cattle by applying a multi-trait test statistic. We detected two QTL at BTA6:90,486,780 and at BTA25:1,455,150 that affect a broad range of economically important traits, including traits related to body conformation and calving, milking speed and longevity. Furthermore, a QTL at BTA6:90,486,780 showed evidence of ongoing selection, whereas the rest of detected QTL showed no or weak evidence of selection. Our findings provide a comprehensive overview of major QTL that segregate in Brown Swiss cattle and revealed pleiotropic effects that may be relevant for future breeding applications.

## Declarations

### Acknowledgements

We acknowledge the Arbeitsgemeinschaft Süddeutscher Rinderzüchter e.V., the Arbeitsgemeinschaft Deutsches Braunvieh, Braunvieh Austria, Tierzuchtforschung Grub, Chair of Animal Breeding of TU München, the Institute of Animal Breeding from Bayerische Landesanstalt fuer Landwirtschaft and ZuchtData EDV Dienstleistungen GmbH for providing genotype and phenotype data for the Austrian and German Brown Swiss populations. We thank the Swiss Brown Swiss breeding association (Braunvieh Schweiz), Arbeitsgemeinschaft Schweizerischer Rinderzuechter and Qualitas AG for providing genotype and phenotype data for the Swiss Brown Swiss population.

### Availability of data and materials

The genotype and phenotype data were provided by breeding organizations (see acknowledgements section) following the execution of material transfer agreements under the condition of strict confidentiality so are not publically available. Contact details of representatives of the breeding organizations are available upon request from Hubert Pausch.

### Author contributions

Analyzed data: Z-HF HP; Wrote the paper: Z-HF HP; Read, revised and approved the final version of the paper: Z-HF HP

### Ethics approval and consent to participate

Not applicable.

### Consent for publication

Not applicable.

## Additional files

### Additional file 1 Table S1

File format: pdf

Description: Number of Brown Swiss bulls for 56 economically important traits and their classification.

### Additional file 2 Figure S1

File format: pdf

Description: Distribution of significant SNPs by number of traits affected for each of 56 economically important traits.

### Additional file 3 Figure S2

File format: pdf

Description: QTL signals detected from milk production, mammary gland morphology and 99-trait meta-analysis on chromosome 6.

### Additional file 4 Table S2

File format: xlsx

Description: Effects of the lead SNPs at QTL detected from the 99-trait meta-analysis across traits.

### Additional file 5 Figure S3

File format: pdf

Description: Effects of the lead SNP of the QTL on chromosome 25 across traits. CH: Swiss population; DEU: Austrian-German population. Effects are t-values estimated from single-trait associations.

### Additional file 6 Figure S4

File format: pdf

Description: Genetic trends for teat thickness and milking speed from year 1980 to 2015.

