## Supplementary material for "Multi-trait meta-analyses reveal 25 quantitative trait loci for economically important traits in Brown Swiss cattle": Table S1

**Table S1 Number of Brown Swiss bulls for 56 economically important traits and their classification.**

| Trait type | Trait name | N of Animal | |
|---|---|---|---|
| | | Swiss | Austrian-German |
| Milk production | Milk yield | 1874 | 2497 |
| | Fat yield | 1874 | 2497 |
| | Protein yield | 1874 | 2497 |
| | Fat percentage | 1874 | 2497 |
| | Protein percentage | 1874 | 2497 |
| Body Size | Stature | 1875 | 2389 |
| | Chest width | 1873 | 2279 |
| | Body Depth | 1874 | 2389 |
| | Angularity | 1871 | 2381 |
| | Rump length | 1874 | 2181 |
| | Rump width | 1874 | 2387 |
| | Rump angle | 1874 | 2389 |
| | Thurl position | 1872 | 1443 |
| | Backline | 1864 | 2387 |
| | Overall frame | 1867 | 2182 |
| | Overall rump | 1850 | 1443 |
| | Final conformation Score | 1861 | 1414 |
| Leg conformation | Hock angularity | 1866 | 2389 |
| | Hock development | 1862 | 2387 |
| | Foot angle | 1850 | 2389 |
| | Hoof height | 1837 | 2386 |
| | Overall leg conformation | 1854 | 2382 |
| Mammary gland morphology | Fore udder length | 1865 | 2387 |
| | Rear udder width | 1863 | 2389 |
| | Rear udder height | 1865 | 2389 |
| | Suspension ligament | 1861 | 2389 |
| | Udder depth | 1872 | 2389 |
| | Fore udder attachment | 1866 | 1384 |
| | Udder balance | 1865 | 1341 |
| | Teat length | 1872 | 2389 |
| | Teat thickness | 1872 | 1369 |
| | Teat placement (front) | 1872 | 2156 |
| | Teat placement (rear) | 1871 | NA |
| | Teat direction (rear) | 1870 | 2387 |
| | Udder cleanness | NA | 2381 |
| | Overall udder score | 1861 | 2384 |

**Table S1 Number of Brown Swiss bulls for 56 economically important traits and their classification (continued).**

| Trait type | Trait name | N of Animal | |
|---|---|---|---|
| | | Swiss | Austrian-German |
| Fertility | Non-return rate (heifer) | 1811 | NA |
| | Non-return rate (cow) | 1836 | NA |
| | Interval from first to last insemination (heifer) | 1801 | NA |
| | Interval from first to last insemination (cow) | 1820 | NA |
| | Days to first service | 1809 | NA |
| Calving | Direct / maternal calving ease | 1632 / 1532 | 2530/2526 |
| | Direct / maternal birth weight | 1639 / 1747 | NA |
| | Direct / maternal gestation length | 1768 / 1776 | NA |
| | Direct live birth | 1068 | NA |
| Growth and carcass quality | Daily net gain | 1470 | 2529 |
| | Carcass percentage | NA | 2460 |
| | Carcass grading | 1450 (CHATX) | 2527 (EUROP) |
| | Daily net gain (calf) | 1527 | NA |
| | Carcass grading (calf) | 1480 (CHATX) | NA |
| | Milking speed | 1846 | 2517 |
| | Longevity | 1805 | 2529 |
| | Persistency | 1868 | 2530 |
| | Somatic cell score | 1861 | 2517 |
