## Supplementary figures and images for "Multi-trait meta-analyses reveal 25 quantitative trait loci for economically important traits in Brown Swiss cattle"

### Figure S1

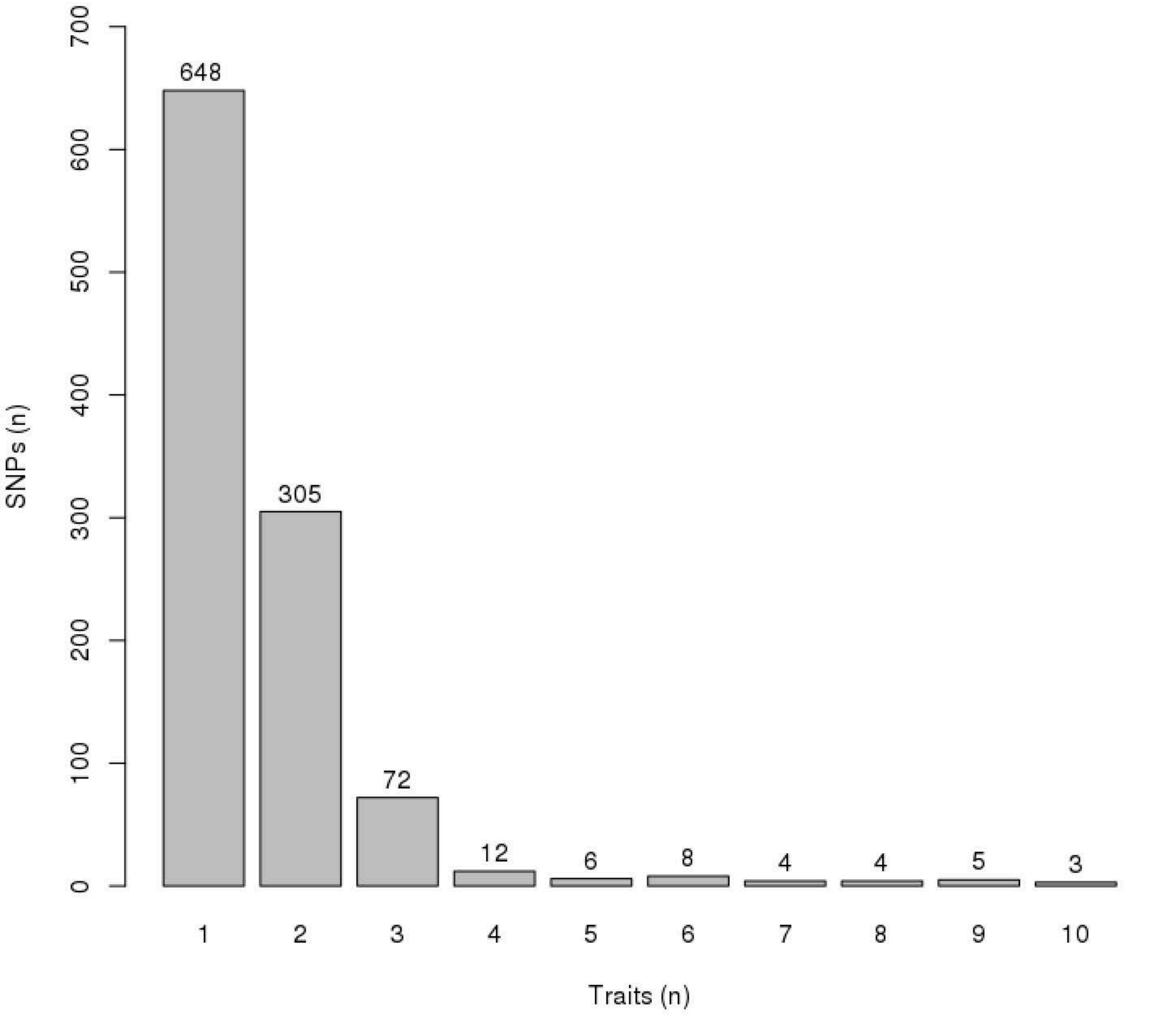

### Figure S2

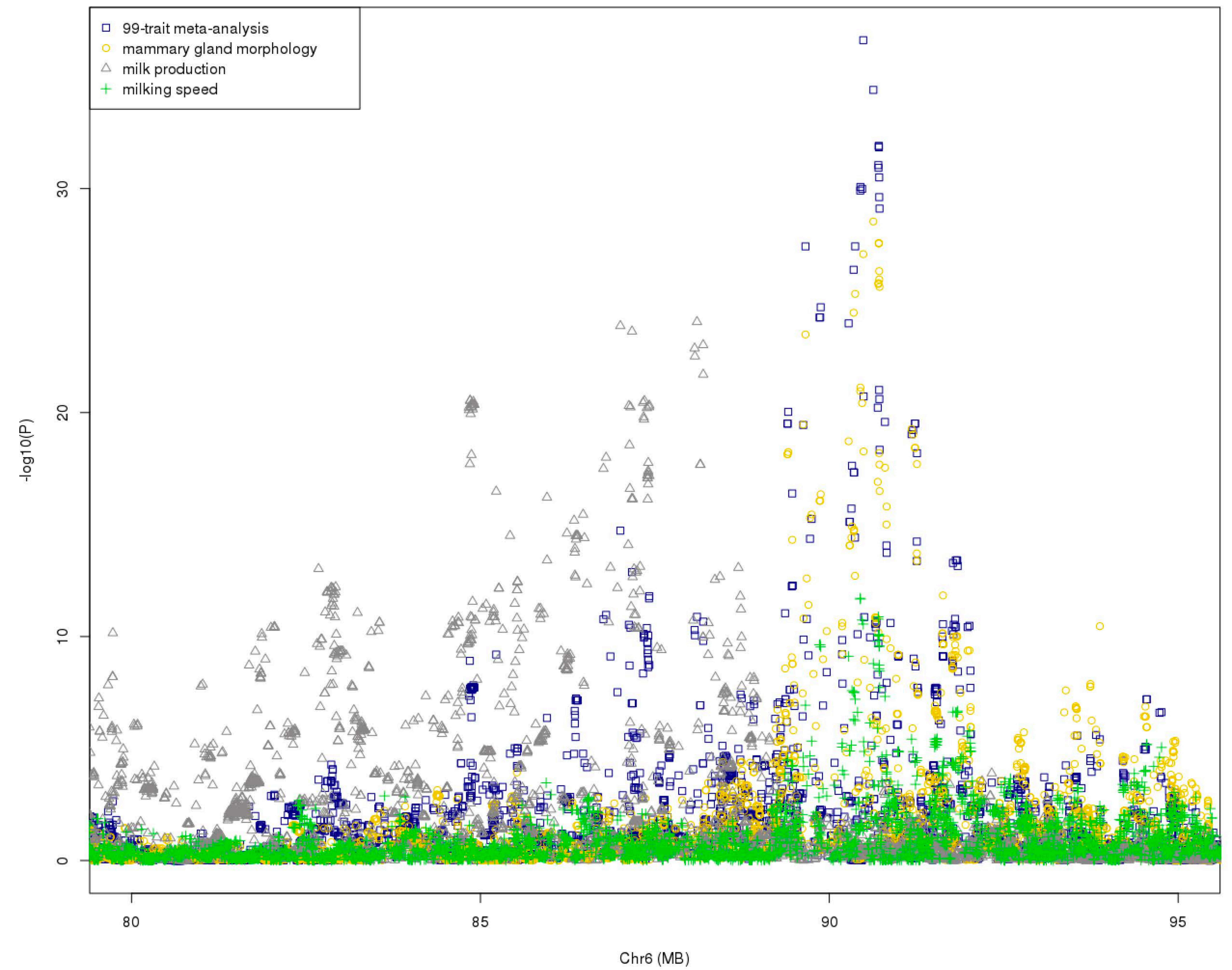

### Figure S3

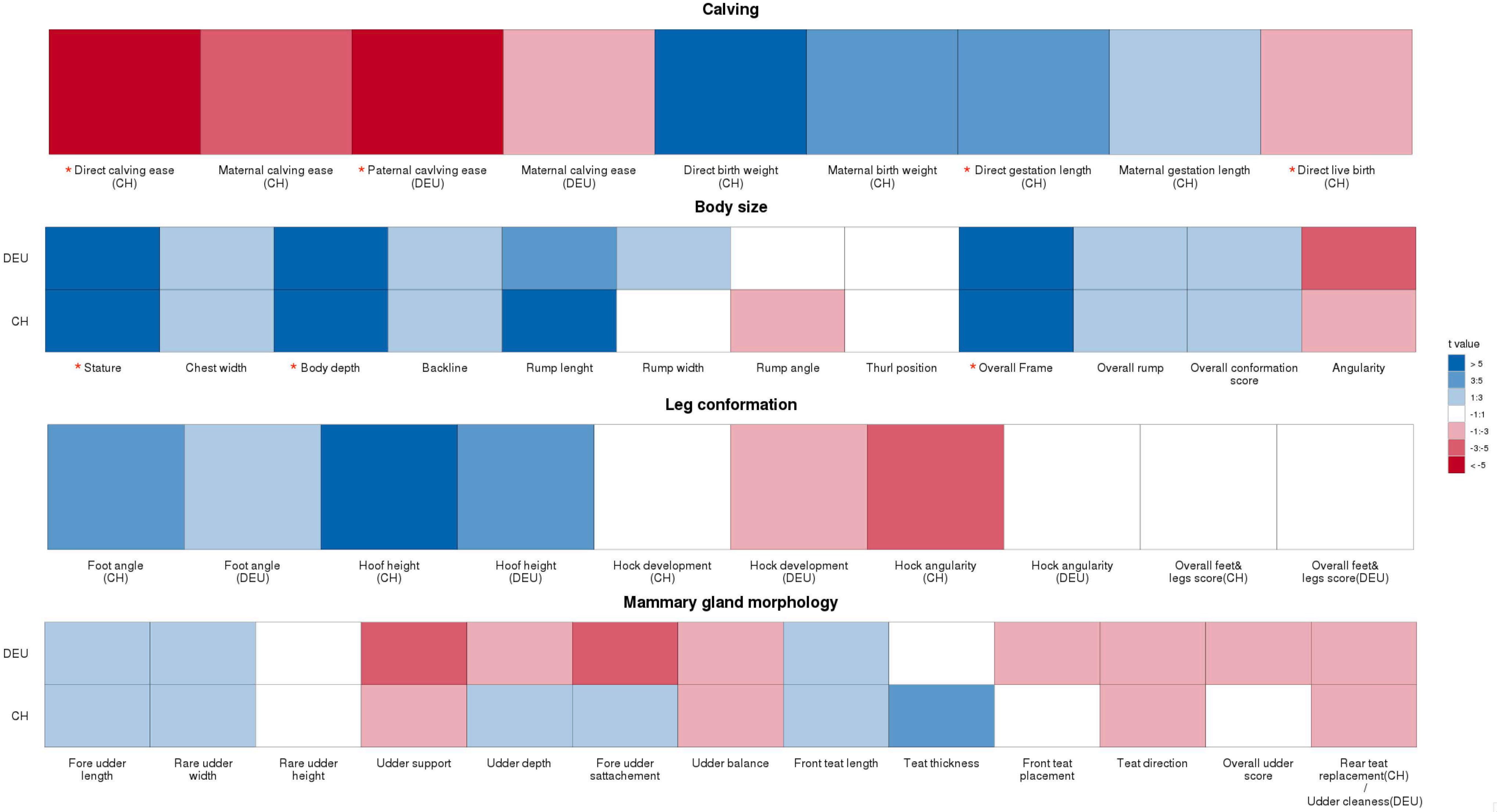

### Figure S4

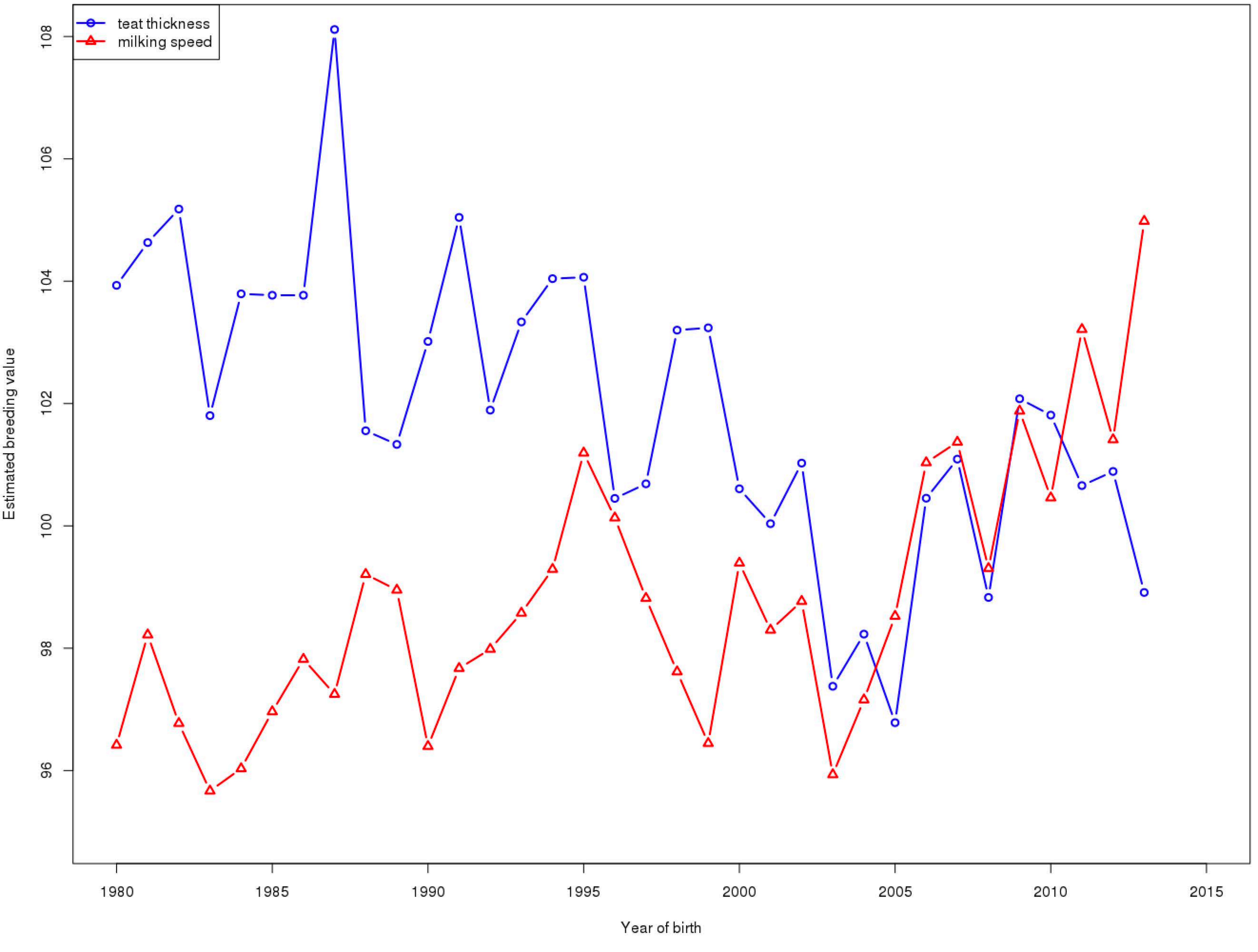
